# Genome-wide transcriptomic response of whole blood to radiation

**DOI:** 10.1101/2025.03.19.644086

**Authors:** Ahmed Salah, Daniel Wollschläger, Maurizio Callari, Heinz Schmidberger, Federico Marini, Sebastian Zahnreich

**Affiliations:** Department of Radiation Oncology and Radiation Therapy, University Medical Center of the Johannes Gutenberg University Mainz, Germany; Institute of Medical Biostatistics, Epidemiology and Informatics (IMBEI), University Medical Center of the Johannes Gutenberg University Mainz, Germany; Fondazione Michelangelo, Milan, Italy; Research Center for Immunotherapy (FZI), Mainz, Germany

**Keywords:** Ionizing radiation, Whole blood, Transcriptomics, RNA-seq, Radiotherapy, Biodosimetry

## Abstract

The hematological system is impacted in nearly all ionizing radiation exposure scenarios. Whole transcriptome data offer detailed insights into blood’s radiation response, crucial for radiotherapy and biodosimetry. We conducted genome-wide RNA-seq analysis on blood from three donors irradiated *ex vivo* with X-rays and incubated for 2h and 6h. Gene expression was subject to strong inter-donor variation and time post-exposure. After 0.5, 1, 2, and 4 Gy X-rays, 5, 33, 84, and 364 genes (2h) and 72, 99, 274, and 607 genes (6h) were differentially expressed (DEG), compared to 0 Gy. The corresponding number of the inferred transcription factors was 255, 253, 274, and 292 after 2h and 214, 245, 262, and 279 after 6h. In sham-irradiated blood, 924 DEGs and 165 transcription factors were affected by *ex vivo* incubation alone. We identified 34 radioresponsive DEGs not previously described, 8 and 9 showing significant positive or negative correlations with dose, respectively, including GPN1, MRM2, G0S2, and PTPRS. DNA damage signaling pathways were affected from the lowest dose, with doses ≥ 2 Gy additionally triggering proinflammatory responses. This first genome-wide RNA-seq study of *ex vivo* X-ray-exposed human blood reveals novel radiosensitive genes, transcription factors, and pathways, enhancing the understanding of the consequences of diagnostic, therapeutic, or accidental exposures on the highly radioresponsive hematological system.

## Introduction

Exposure to ionizing radiation (IR) *in vivo* commonly results in blood irradiation, representing one of the most radioresponsive and -sensitive tissues (1). The hematological system’s cellular stress response to radiation and its impact on systemic homeostasis is gaining attention due to its relevance in radiotherapy (RT) for benign or malignant conditions and its use as a biodosimeter in radiation accidents (2,3).

IR is a potent inducer of DNA damage, activating a complex DNA damage response (DDR) primarily mediated by the kinases ataxia telangiectasia mutated (ATM) and Rad3 related (ATR) or DNA-dependent protein kinase catalytic subunit (DNA-PKcs) (4,5). These kinases target key transcription factors such as p53, NF-kappaB, BRCA1, and AP-1, driving cellular outcomes like transient or permanent cell cycle arrest (including premature senescence), apoptosis, and immunogenic signaling (5–7).

Hematologic cells, particularly lymphocytes, are highly sensitive to apoptosis and exhibit immunogenic responses even at low doses of IR (8,9). These reactions enhance the efficacy of low-dose radiotherapy (RT) for benign degenerative and inflammatory conditions such as osteoarthritis, epicondylitis, scapulohumeral periarthritis, and heel spur (9). Around 0.5 Gy, lymphocytes display hypersensitivity and a non-linear dose-response, which is leveraged in anti-inflammatory RT regimens for such pathologies (10). Conversely, in RT for malignancies involving tumor doses of 2 Gy or more per fraction, these effects contribute to leukopenia and hematopoiesis suppression, representing negative prognostic factors in many solid tumor entities (11).

Recent attention has shifted to combining RT with immuno-oncological antitumor strategies, where lymphocytes mediate local and systemic (abscopal) effects through immunogenic tumor cell death and signaling (12). However, a lack of synergy in these combinations is often due to RT’s adverse impact on the hematological system (13). The depletion of immune cells induced by IR, along with their release of cytokines and danger signals, exerts immunomodulatory effects on both the local microenvironment and systemic responses (14). These effects influence the therapeutic efficacy of RT and contribute to radiotoxicity in normal solid tissues (14,15).

Beyond quantifying immune cell populations, DNA damage, and metabolic or protein markers, the analysis of gene expression and downstream signaling pathways provides the most detailed insights into the global cellular responses of the hematologic system to IR (16,17). Gene expression studies on human whole blood or peripheral blood mononuclear cells (PBMCs), irradiated *ex vivo* or *in vivo*, have primarily been used in biodosimetry to identify exposed individuals, estimate absorbed doses, and predict individual responses (18).

These studies used primarily microarrays and quantitative polymerase chain reaction (qPCR) to analyze gene expression in irradiated blood cells across varying doses, dose rates, radiation qualities, and time points (18). A panel of IR-responsive genes exhibiting time- and dose-dependent transcriptional regulation have been identified, including FDXR, APOBEC3H, CCNG1, PHPT1, MDM2, BBC3, and CDKN1A (18). Recent studies have shifted focus from general DDR to understanding the IR-associated transcriptional inflammatory signature in the blood (3,16,19–21). This knowledge could enhance therapeutic outcomes for cancer patients in multimodal radiation immuno-oncology and mitigate RT-associated side effects. However, genome-wide transcriptional analyses of blood cells in response to radiation by RNA-seq remain limited. Notably, only one study has employed third-generation long-read Oxford Nanopore Technologies (ONT) RNA-seq, focusing on isolated PBMCs exposed to a single 2 Gy dose and analyzed 24h post-irradiation (22). Compared to microarray analysis, RNA-seq offers rapid, in-depth profiling of the genome-wide transcriptome, enabling the detection of novel transcripts, a broader dynamic range, greater specificity and sensitivity, and the identification of rare transcripts (23). To better understand the IR-associated genome-wide transcriptome response in whole blood, RNA-seq approaches are crucial. Further studies are needed to develop an integrative understanding of the effects and consequences of diagnostic, therapeutic, and accidental exposures on the highly IR-responsive hematological system.

To identify novel gene signatures and signaling pathways of IR exposure in blood, we conducted genome-wide transcriptome analyses of whole blood from three healthy donors exposed to 0, 0.5, 1, 2, and 4 Gy of X-rays, with samples collected 2h and 6h post-irradiation using short-read Illumina RNA-seq.

## Materials and Methods

### Blood sampling, irradiation, culturing, and RNA isolation

Whole blood was collected from three healthy donors (two males and one female) by venipuncture into EDTA tubes (S-Monovette® EDTA K3E, Sarstedt, Nuembrecht, Germany). Ethical approval was obtained from the Medical Association of Rhineland-Palatinate [No. 2023-17191], and all research was performed in accordance with relevant guidelines and regulations. All donors provided informed consent and research has been performed in accordance with the Declaration of Helsinki. X-ray irradiation was conducted using a D3150 X-Ray Therapy System (Gulmay Ltd., Surrey, UK) at 140 kV with a dose rate of 3.6 Gy/min at room temperature. Samples were exposed to 0.5, 1, 2, and 4 Gy X-rays, with sham-irradiated (0 Gy) controls maintained under identical conditions in the control room. After irradiation, 1 ml of whole blood per sample was mixed with 1 ml preheated (37 °C) X-VIVO™ 15 media (Lonza Group Ltd., Basel, Switzerland) containing 10% heat-inactivated fetal calf serum (Bio&SELL GmbH, Feucht, Germany) (56°C, 30 min). The cell suspension was cultured in 6-well plates (Greiner Bio-One, Kremsmünster, Austria) and incubated at 37 °C, 5% CO2, and a humidified atmosphere for 2h or 6h. RNA was extracted using the QIAamp RNA Blood Mini Kit (Qiagen, Hilden, Germany) according to the manufacturer’s protocol.

### RNA sequencing

RNA library preparation and transcriptome sequencing from 30 samples were conducted by StarSEQ GmbH (Mainz, Germany). Globin RNA and rRNA were depleted using the NEBNext Globin & rRNA Depletion Kit (New England Biolabs, Ipswich, USA). RNA quality was assessed with a Bioanalyzer and Qubit. Libraries were prepared with the NEBNext UltraExpress Directional RNA Library Preparation Kit. Sequencing was performed on a NextSeq 2000 instrument (Illumina, San Diego, CA, USA) utilizing a P3 flow cell and XLEAP-SBS chemistry, generating paired-end reads of 150 nucleotides. Base calling was executed using NextSeq 1000/2000 Control Software Suite v1.7.1, and the data was converted into FASTQ format with DRAGEN BCL Convert version 4.2.7.

### Processing of RNA-seq data

Quality control on the sequencing data was performed with the FastQC tool (version 0.12.1., https://www.bioinformatics.babraham.ac.uk/projects/fastqc/).

Transcript abundance was estimated using Salmon (version 1.10.3) (24) with a decoy-aware transcriptome index based on GENCODE version 45. The results were then summarized to the gene level using the tximeta R package (version 1.22.1) (25).

Principal Component Analysis (PCA) plots were performed using the pcaExplorer package (26,27) including the top 500 most variable genes.

Differential expression analysis was conducted using the DESeq2 package (version 1.44.0) (28), with the false discovery rate (FDR) cutoff set to 0.05. Sham-irradiated samples were used as the reference level. The DESeq2 statistical workflow accounted for dose effects and inter-donor variability. Genes from the 0.5, 1, 2, and 4 Gy groups were considered differentially expressed if their adjusted p-value (Benjamini-Hochberg procedure) was less than 0.05.

Log2 fold change effect sizes were estimated using the apeglm shrinkage estimator (version 1.26.1) (29). A time-series analysis workflow from DESeq2 was applied by specifying interaction terms for dose and time points, enabling a comparison of radiation effects at 2h *versus* 6h post-irradiation.

Gene enrichment analysis of the differentially expressed genes (DEGs) was conducted using ClusterProfiler (version 4.12.6) (30–32), with all expressed genes used as the background dataset. The enrichment results were visualized and summarized using the GeneTonic package (version 2.8.0) (26,33).

Gene expression profiles were displayed as heatmaps, with color-coded standardized Z-scores for expression values. These values were obtained after regularized logarithm transformation, facilitating comparison across samples.

### Transcription factors activity inference and deconvolution analysis

The transcription factor (TF) inference score was calculated using a Univariate Linear Model implemented in the Decoupler package (version 2.10.0) (34). Immune cell relative abundance was estimated with Omnideconv (version 0.1.0) (35) employing the BayesPrism method (36). A blood single-cell dataset from Tabula Sapiens (TS_Blood.h5ad retrieved from https://doi.org/10.6084/m9.figshare.14267219.v5), filtered to include a minimum of 200 and a maximum of 500 cells per cell type, was used as the reference for these analyses.

### Gene set variation analysis

Gene Set Variation Analysis (GSVA) was performed with the GSVA package (version 1.52.3) (37). Gene Ontology (GO) IDs were obtained from the org.Hs.eg.db package (version 3.19.1). The resulting GSVA score matrix was used for differential pathway-level expression analysis, which was conducted using the limma package (version 3.60.6) (38).

### Meta-analysis of within-donor Pearson correlation

The correlation between normalized gene counts and radiation dose was calculated while accounting for repeated measures from each donor, which introduced dependencies in the data. Donor-specific correlations were computed (39) and then pooled using a fixed-effects meta-analysis based on Rubin’s rules (40). The meta-analysis provided the estimated correlation, p-value, and 95% confidence interval, which were used to identify radiosensitive genes.

## Results

### Data exploration and differential expression analysis

Whole blood from three healthy donors was exposed to X-rays at graded doses (0, 0.5, 1, 2, and 4 Gy) and incubated at 37 °C for 2h and 6h to mimic *in vivo* gene expression dynamics. Total RNA was isolated from the samples and sequenced using 150 bp short reads, with an average of 10 million aligned reads per sample (Supplementary Fig. S1).

Exploratory data analysis showed no gene expression grouping by radiation dose but identified clear donor-related clusters strongly influenced by sex and incubation time. PCA further confirmed that donor and time effects accounted for the greatest variability (Fig. 1A). Samples from different donors formed distinct clusters at both time points, with male donor clusters being closer to each other than to the female donor cluster. Differentially expressed genes (DEGs) were calculated, accounting for the donor and time effects in the analysis design. The number of significantly up- and downregulated genes was dependent on both the dose and the time after irradiation (Fig. 1B and C).

**Figure 1:**
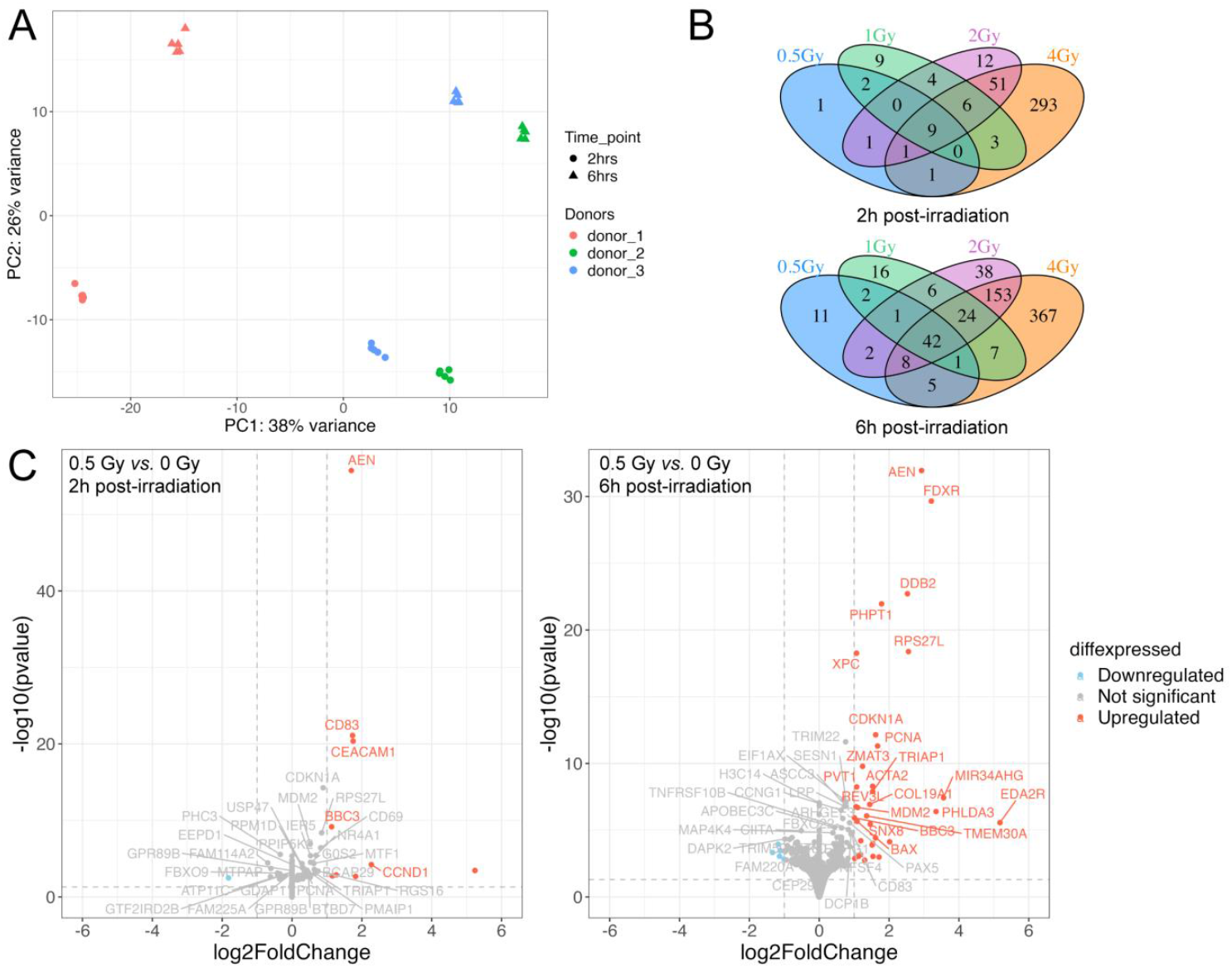
Gene expression analysis of *ex vivo* irradiated whole blood from 3 healthy donors by short-read RNA-seq 2h and 6h post-exposure. (A) Principal Component Analysis (PCA) plot showing the variability due to donor (PC1) and time (PC2) effect. Each data point for a donor represents a radiation dose. (B) Venn diagrams providing the number of DEGs with different doses at 2h and 6h post-exposure. (C) Volcano plots representing the effect on gene expression levels of 0.5 Gy compared to 0 Gy X-rays at 2h and 6h after exposure.

2h post-irradiation, differential gene expression analysis revealed 15, 33, 84, and 364 DEGs at 0.5, 1, 2, and 4 Gy, respectively, compared to 0 Gy. AEN, CD83, CDKN1A, BBC3, RPS27L, MDM2, CD69, NR4A1, and IER5 were consistently upregulated across all radiation doses, confirming their role as main players in conserved mechanisms of IR response. Dose-specific regulation was observed for the upregulation of CEACAM1 after 0.5 Gy and the upregulation of HMGN2p5 and TNFRSF10B as well as the downregulation of DHFRP1, LINC00472, and RN7SL731P after 1 Gy. After exposure to 2 Gy and 4 Gy, 10 and 281 unique DEGs were found, respectively. The full list of DEGs, including dose- and time-specific changes *versus* sham-irradiated controls, is provided in Supplementary Table S1.

6h post-irradiation, differential gene expression analysis identified 72, 99, 274, and 607 DEGs at 0.5, 1, 2, and 4 Gy, respectively, compared to 0 Gy. 41 genes, including FDXR, AEN, XPC, BBC3, PAX5, APOBEC3C, DDB2, and BAX, were consistently differentially expressed across all radiation doses. Unique DEGs at specific doses included H3C14, GNL3LP1, and RHEBP2 after 0.5 Gy, BRCA2, SOCS4, SERGEF and RNF141 after 1 Gy, NOTCH2, NOLC1, PIBF1 and CCAR2 after 2 Gy and DDX21, BID, MXD4 and VEGFA after 4 Gy. These findings underscore both generalized and dose-specific transcriptional responses of whole blood at the two investigated time points post-exposure, highlighting the dynamic and context-dependent nature of gene expression changes in response to IR.

Next, we analyzed the dose- and time-dependent expression of IR-responsive genes in greater detail. This investigation also aimed to identify new radiosensitive genes in human whole blood using a genome-wide RNA-seq approach. Table 1 presents correlation analysis data for selected well-known and all newly identified genes as a function of dose at 6h post-irradiation, along with information on their significant expression compared to sham-irradiated controls 2h and 6h after exposure. Examples of dose-dependent gene expression are shown in Fig. 2A. We identified 17 genes with a significant correlation and high correlation coefficients that, to our knowledge, have not been previously described as radioresponsive in *ex vivo* photon-irradiated human blood.

**Table 1:**
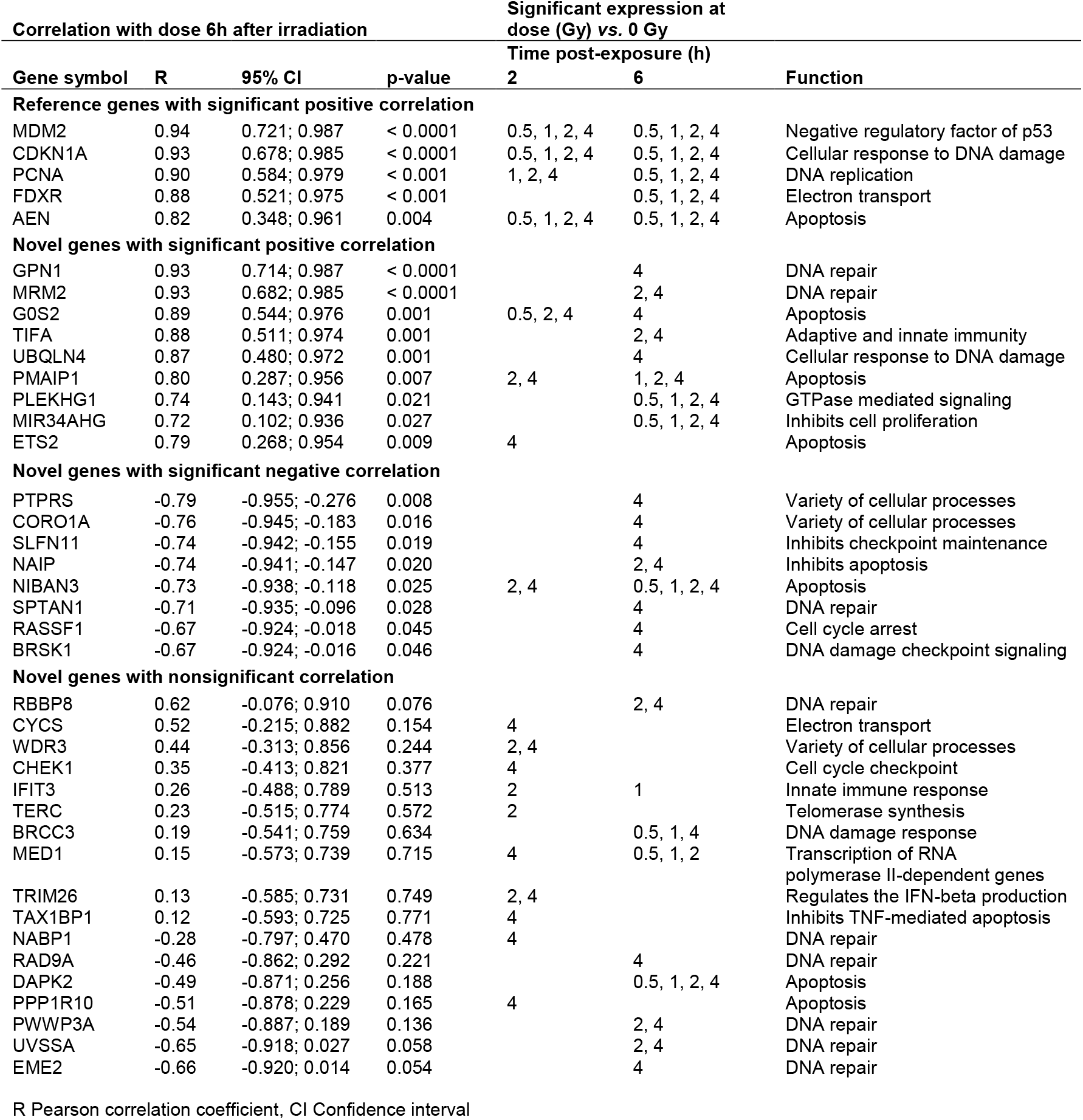
Meta-analysis of within-donor Pearson correlation of selected known and in this study newly identified radioresponsive genes in human whole blood with applied dose 6h after irradiation and their significant expression at a given dose compared to the sham-irradiated sample 2h and 6h after irradiation.

**Figure 2.**
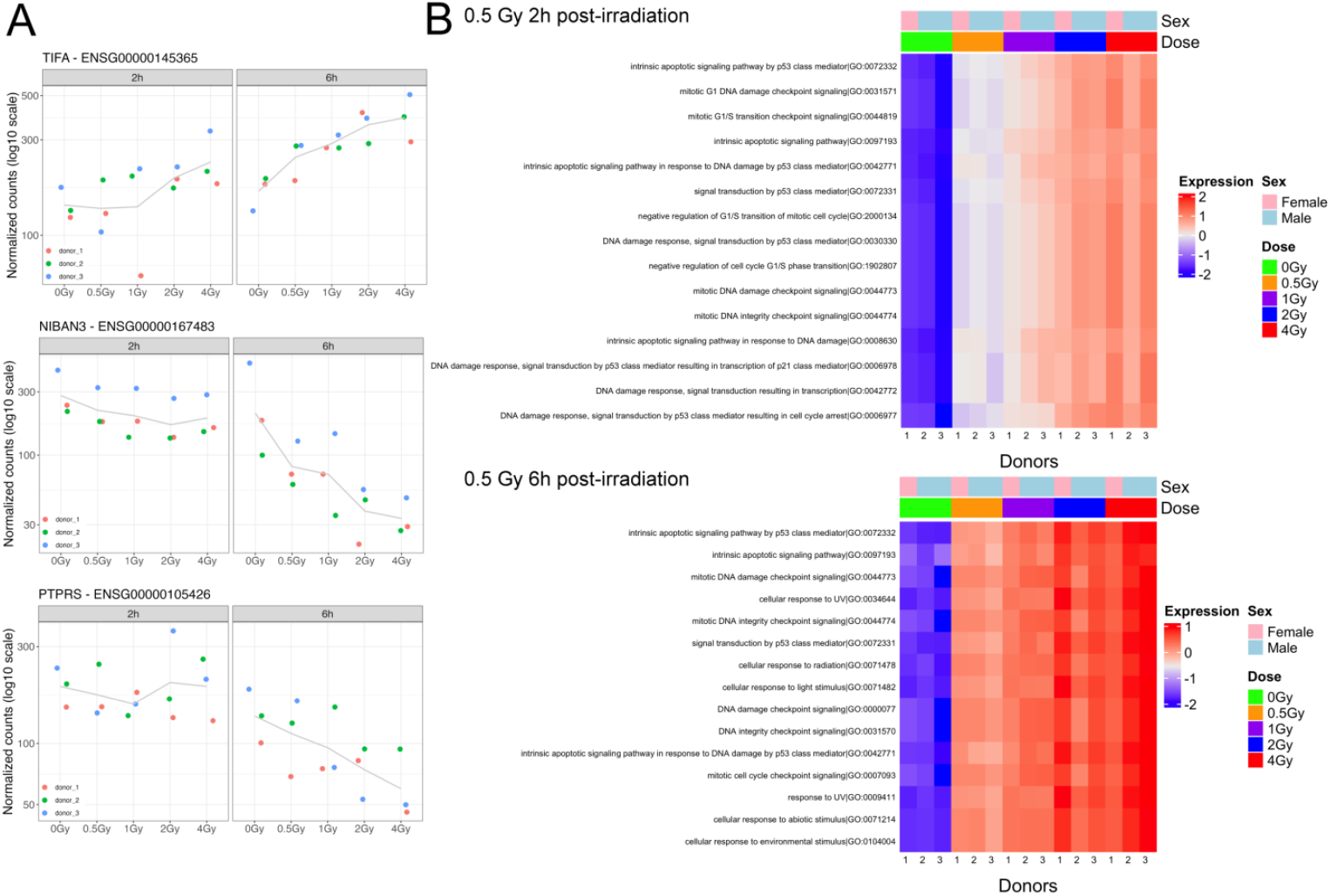
(A) Exemplary significant dose-response relationships for 3 newly identified radioresponsive genes in *ex vivo* human blood 2h and 6h after X-ray irradiation. (B) Heatmaps of the regulation of the most enriched signaling pathways in response to a low X-ray dose of 0.5 Gy compared to 0 Gy 2h and 6h after irradiation.

Eight genes exhibited a significant negative correlation with dose: PTPRS, CORO1A, SLFN11, NAIP, NIBAN3, SPTAN1, RASSF1, and BRSK1. Another nine genes showed a significant positive correlation with dose: GPN1, MRM2, G0S2, TIFA, UBQLN4, PMAIP1, PLEKHG1, MIR34AHG, and ETS2. Compared to established radioresponsive genes such as CDKN1A and MDM2, most newly identified genes showed significant differential expression at higher doses 6h post-irradiation. Only PLEKHG1, MIR34AHG, and NIBAN3 exhibited significant regulation across the entire dose range.

To investigate pathway activity regulation, we conducted a functional enrichment analysis on the DEGs. Even at low radiation doses from 0.5 Gy, the DNA damage and cell cycle-regulating signaling pathways were the most strongly and dose-dependently enriched at both time points. These results are depicted in Fig. 2B, emphasizing the early and robust activation of these critical pathways in response to radiation exposure.

To explore a specific DDR pathway in greater detail, we focused on the “intrinsic apoptotic signaling pathway in response to DNA damage” (GO:0008630) as an example. Genes activating this pathway, identified among our DEGs, demonstrated clear dose-response activation, as illustrated in Supplementary Fig. S2. This highlights the pathway’s pivotal role in the cellular response to IR-induced DNA damage.

### Radiation impact on the inflammatory response

We next examined the effect of IR on the transcriptomic inflammatory response, primarily observed at radiation doses ≥ 2 Gy. Key inflammatory DEGs significantly stimulated at these doses included CCL4, NFKB1, IL1B, CD70, CD83, CCL4, CXCL8, and TNF (Fig. 3A). Signaling pathways associated with the immune response, particularly pro-inflammatory signaling, showed increased and dose-dependent enrichment at doses ≥ 2 Gy at both time points (Fig. 3B and C). These findings indicate a significant role of pro-inflammatory activity in the transcriptomic response to higher radiation doses. Certain inflammatory genes, such as TNF, PTGS2, and IL1B, decreased from their high initial expression at 6h post-irradiation. To determine whether the immune response had diminished, we analyzed how DEGs from the 4 Gy sample at 6h clustered at the pathway level. Functional enrichment analysis revealed that pro-inflammatory pathways remained upregulated at 6h (Supplementary Table S2), suggesting that while individual gene expression levels may decline over time, the overall activation of inflammatory pathways persists.

**Figure 3.**
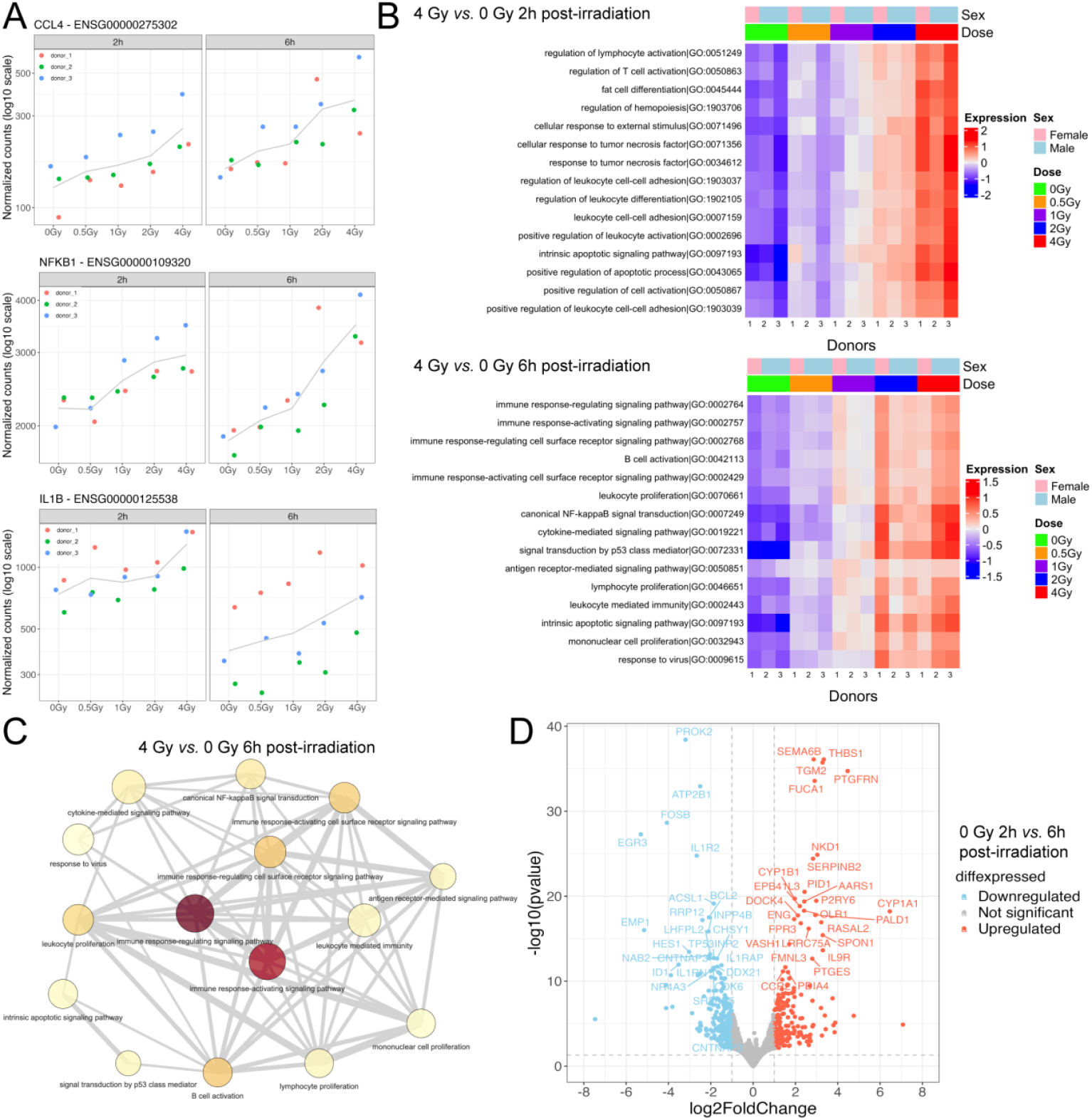
Radiation effects on the inflammatory response in whole blood. (A) Dose-response relationships for selected inflammatory genes. (B) Heatmaps showing the 15 most enriched inflammatory signaling pathways 2h and 6h after irradiation with 4 Gy. (C) Network plot of the top enriched immunogenic pathways after 4 Gy irradiation and 6h of incubation. The node color represents the degree of significance (from light yellow to dark red) by adjusted p-value and grey lines indicate the presence of shared genes between each connected node. (D) Volcano plot for the effect of ‘time in culture’ on differentially expressed genes in sham-irradiated samples at 2h vs. 6h after the start of blood incubation.

To investigate the change in the degree of regulation in some inflammatory gene expressions in response to time, we conducted a differential expression analysis comparing the sham-irradiated group at each time point (0 Gy at 2h *versus* 0 Gy at 6h). This analysis identified 924 DEGs stimulated solely by the duration of *ex vivo* blood incubation (Fig. 3D and Supplementary Table S3). Gene Set Enrichment Analysis (GSEA) of these 924 DEGs revealed that the enriched pathways were predominantly associated with anti-inflammatory activity (Supplementary Fig. S3, Supplementary Table S3). These findings suggest that the waning of specific inflammatory gene responses over time may be influenced by anti-inflammatory processes activated during prolonged blood incubation.

To account for the time-dependent effects of *ex vivo* blood incubation on gene expression, we refined our analysis to focus exclusively on the late effects of IR on DEGs. The number of DEGs identified for 2 Gy and 4 Gy comparisons at 6h, relative to their earlier time points, was 42 and 74, respectively. These genes were involved in the DDR and the immune response, indicating that the late effects of higher radiation doses still elicit immune reactions.

We utilized GSVA to perform differential expression analysis at the pathway level, aiming to corroborate enrichment results when comparing early to late effects of IR. No differentially regulated genesets were detected at a p-value threshold of 0.05, indicating no significant change between late and early effect after accounting for time-related confounding changes in blood by *ex vivo* incubation.

### Quatificaion of radiation impact on transcription factors

To further investigate the upstream regulation of the transcriptional radiation response of blood, we inferred TFs activity using a univariate linear model with the decoupleR R package. There was a considerable transcriptional response in whole blood involving TFs after IR exposure. However, in contrast to the number of DEGs, the number of significantly regulated TFs was largely independent of dose and time post-exposure. Compared to sham-irradiated samples, the number of TFs significantly regulated following irradiation with 0.5, 1, 2, and 4 Gy was 255, 253, 274, and 292 after 2h and 214, 245, 262, and 279 after 6h, respectively. However, as shown in Fig. 4A, a dose-dependent upregulation was still observed for the top 25 variable TFs 2h post-irradiation, followed by a downregulation at 6 h. An exception to this pattern was AHRR, a TF known to induce immune activity, which showed general upregulation 6h after irradiation. A full list of TFs, including dose- and time-specific changes *versus* sham-irradiated controls, is provided in Supplementary Table S5.

**Figure 4.**
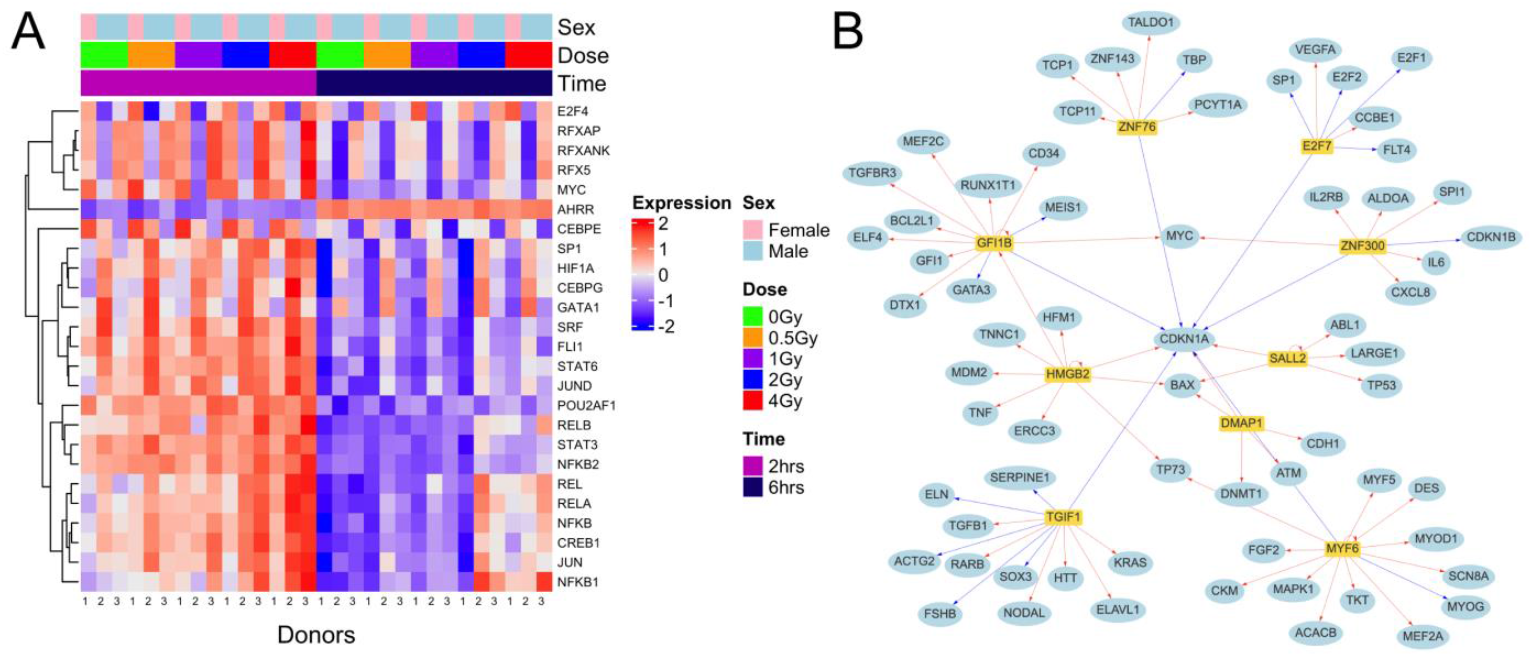
Analysis of transcription factors in irradiated whole blood. (A) Heatmap of the top transcription factors regulated by ionizing radiation. (B) Network plot for the radioresponsive transcription factors (yellow rectangles) affecting target genes (blue circles) including CDKN1A. Blue and red lines represent downregulation and upregulation, respectively.

Among the top 25 significantly regulated TFs in response to all radiation doses and time points compared to sham-irradiated cells, DMAP1, E2F7, GFI1B, HMGB2, MYF6, SALL2, TGIF1, ZNF300, ZNF76 were found to be commonly shared. E2F7, GFT1B, MYF6, TG1F1, ZNF300, and ZNF76 were consistently downregulated and DAMP1, HMGB2, and SALL2 were consistently upregulated. Interestingly, inferred TF activities showed that downregulated TFs correlated with CDKN1A downregulation, while upregulated TFs were linked to its upregulation in response to IR-induced DNA damage. The relationship between TFs regulating CDKN1A and their target genes is depicted in Fig. 4B.

Since we observed a time effect due to *ex vivo* blood incubation on DEGs in sham-irradiated samples, we also examined this effect for TFs. Our analysis identified 165 TFs influenced by the time effect of *ex vivo* incubation, with 128 being downregulated (Supplementary Table S6). This explains the general downregulation pattern of TFs observed in the heatmap for samples incubated for 6h (Fig. 4A). Similar to the DEGs, the affected and predominantly downregulated TFs were associated with the inflammatory response. Together, TF activity highlighted the regulation of TFs affecting inflammation and DDR after IR exposure, influenced by the *ex vivo* incubation effect.

## Discussion

Exposure to IR is associated with acute and late health effects, with inflammatory reactions playing a critical role in radiation accidents and clinical applications. The hematologic system, almost always affected in IR exposure scenarios, is highly susceptible to radiation effects. It is therefore involved in many systemic radiation responses and utilized in biodosimetric assays. Understanding the blood’s response to IR is increasingly significant for biological dosimetry following radiation accidents and planned therapeutic or diagnostic exposures. Transcriptome analysis offers the most comprehensive view of global cellular responses to IR, and various studies have identified multiple IR markers of blood gene expression (41,42). To our knowledge, this study represents the first genome-wide transcriptomic short-read RNA-seq analysis of *ex vivo* photon-irradiated human whole blood. Blood samples from three healthy donors were exposed to X-rays with a range of doses, followed by RNA extraction and sequencing after 2h and 6h of *ex vivo* incubation. Our analysis focused on identifying differentially expressed genes, functional annotation using GO-based pathways, estimating transcription factor inference activity, and immune cell population deconvolution.

Our initial exploratory data analysis revealed that the sex difference among the three donors contributed to the highest variability in the data. This variation was noticeable at the level of both expressed genes and TFs. However, given the small sample size, this effect is more appropriately attributed to inter-donor variation rather than being conclusively sex-specific. The well-known inter-individual variability in gene expression at baseline and after irradiation has been attributed to various factors, including age, sex, smoking status, body mass index, and genetic factors. Agbenyegah *et al*. (43) investigated the baseline expression of 10 radiosensitive genes, including FDXR, DDB2, WNT3, and CD177, in 200 healthy male and female donors. They found significant associations between gene expression and factors like sex (e.g., DDB2), age (e.g., CD177), or both sex and age (e.g., STAT4). Similarly, Li *et al*. (44) observed that IR-induced expression changes in highly radioresponsive genes such as CDKN1A, PCNA, TNFSF4, POLH, and CCNG1 were significantly greater in female blood samples compared to male samples at γ-radiation doses of 0.5 to 8 Gy, 24h post-exposure. However, many gene expression patterns are also influenced by genetic factors. Correa and Cheung (44) demonstrated that FDXR and CDKN1A expression after irradiation in lymphoblastoid cells was significantly more similar within monozygotic twin pairs than between twin pairs, emphasizing the role of genetic background. We also detected a striking donor difference in a deconvolution analysis of the RNA-seq data to estimate the proportions of different immune cell populations in the blood samples (45). The individual heterogeneity in blood cell composition has been described as another significant factor contributing to variability in blood gene expression profiles (46,47). Notably, in this study, the female donor consistently exhibited a higher fraction of neutrophils and a lower fraction of lymphocytes compared to the two male donors (Supplementary Fig. S4). This donor-dependent variation in leukocyte populations could therefore be the main factor for the observed strong differences in gene expression and TF values between the sexes. Importantly, neither irradiation nor *ex vivo* incubation altered the immune cell composition across samples, as confirmed by the deconvolution analysis. Inter-individual variance in gene expression is a well-documented and critical factor in biodosimetry after accidental IR exposures, where pre-exposure samples are usually unavailable for distinguishing non-exposed from exposed individuals (21,43,48,49). For example, Paul *et al*. (48) used the Agilent one-color microarray workflow with a Nearest Centroid Classifier and leave-one-out cross-validation to identify IR-induced gene expression patterns without pre-exposure samples. For this reason, genes with low inter-individual variability, such as FDXR, are usually selected in biodosimetry to identify exposed individuals and to estimate absorbed doses based on *ex vivo* dose-response relationships (49). Consequently, whole-genome transcriptome screenings, as conducted in this study, are essential to discovering new radioresponsive genes with low interindividual variability and expanding dose ranges for biodosimetric purposes.

As shown by PCA, sampling time was another factor significantly influencing gene expression. This was observed in irradiated and sham-irradiated blood samples incubated *ex vivo* for 2h or 6h. Longer incubation times resulted in more dose-dependent DEGs, greater regulation of these genes, and activation of more associated signaling pathways. The overall classes of biological functions affected remained unchanged over time. These findings align with Paul *et al*. (48), who demonstrated that a classifier based on a single 74-gene set could accurately predict absorbed radiation doses over an *ex vivo* incubation period of 6–24 h post-irradiation. Thus, dose-dependent gene expression signatures are considered temporally stable from about 6h post-irradiation, making them suitable for retrospective biodosimetric applications. Additionally, 10 groups of DDR genesets were time-dependently regulated across all radiation doses, with increasing dose-dependent intensity indicated by their adjusted p-values in the present study. These commonly regulated pathways included the intrinsic apoptotic pathway (GO:0097193), signal transduction by a p53 class mediator (GO:0072331), and mitotic DNA damage checkpoint signaling (GO:0044773). These findings highlight the dynamic regulation of DDR pathways and their increasing activation in response to IR over time. However,we showed that *ex vivo* incubation significantly influenced gene expression already in sham-irradiated controls. The ‘time in culture effect’ on blood gene expression is a well-recognized interfering factor (47,48). Our differential expression analysis of sham-irradiated samples identified the regulation of more than 900 DEGs between 2h and 6h of blood incubation. This incubation effect also impacted the regulated genes in irradiated samples and shifted the direction of associated signaling pathways, particularly those related to inflammation. GSEA revealed that inflammatory pathways were downregulated, with a negative Normalized Enrichment Score. This could explain the observed decrement of the inflammatory response at 6h compared to 2h post-irradiation.

We incorporated the time effect into the statistical design to account for confounding factors and accurately interpret the IR-associated transcriptomic response. Accounting for this temporal effect revealed that most significantly IR-induced genes were linked to the DDR. Key genes, such as FDXR, PHPT1, ZMAT3, PVT1, RPS27L, ACTA2, DDB2, AEN, PCNA, and XPC, were significantly upregulated in response to X-ray doses ranging from 0.5 Gy to 4 Gy at 6h compared to 2h post-exposure. Generally, our findings align well with prior studies identifying radiosensitive genes in blood for biodosimetry, including FDXR, TIGAR, MYC, GADD45A, PLK2, and SESN1 (48,50,51). These results further support the utility of these genes for accurate dose estimation in accidental radiation exposure scenarios. However, previous gene expression biodosimetry studies have predominantly relied on microarrays or qPCR, which offer significantly lower resolution than RNA-seq utilized in this study. Additionally, considerable variability exists between studies due to differences in parameters such as analyzing irradiated whole blood or isolated PBMCs, *ex vivo* or *in vivo* exposure scenarios, species, radiation qualities, or dose rates.

To evaluate the resolution and performance of RNA-seq compared to microarray techniques, we selected studies that analyzed human whole blood and adhered to comparable criteria, including *ex vivo* incubation time, radiation dose, and quality. Differences in experimental conditions and statistical models for extracting DEGs also influence the comparability of studies. For example, microarray-based profiling of whole blood gene expression upon exposure to 2 Gy or 4 Gy X-rays after 2h by Kabacik *et al*. (52) resulted in 296 upregulated genes or 234 upregulated and 1 downregulated genes, respectively. The number of IR-regulated genes thus appeared to be very similar for both doses after 2h, and there was no fundamental difference even after 24h of incubation. In contrast, in RNA-seq, we observed a dose-dependent increase in IR-associated DEGs at both time points with a further effect of time on gene regulation. In a direct comparison 2h after exposure, 62 upregulated and 22 downregulated genes were detected after 2 Gy and 277 upregulated and 87 downregulated genes after 4 Gy. These results emphasize the higher resolution of RNA-seq compared to microarrays and enable the detection of a broader spectrum of IR-associated gene expression changes to expand the panel of potential biomarkers.

The systematic review by Lacombe *et al*. (41) analyzed 24 studies using microarray approaches to measure gene expression in photon-irradiated blood. The authors identified 31 genes significantly altered after at least one radiation dose in 12 studies, including ACTA2, AEN, FDXR, RPS27L, MYC, and CDKN1A. In this study, we confirmed and validated the regulation of these 31 genes in the same directions following X-ray exposure. Moreover, we demonstrated, at least to our knowledge, the significant regulation of several new radiosensitive DDR genes following photon-irradiation of human whole blood, marking a valuable contribution to the field. Among these, 17 genes exhibited a significant positive or negative correlation with radiation dose 6h post-exposure. For example, novel genes such as GPN1 and MRM2 displayed strong and highly significant dose correlations but showed significantly increased expression levels only at higher doses (≥ 2 Gy) after 6h. In contrast, well-known radioresponsive genes like MDM2 and CDKN1A, with similar positive dose correlations, were significantly upregulated across the entire dose range (0.5–4 Gy) and at both analyzed time points. Future research should focus on exploring and validating the potential of these newly identified radiosensitive genes for applications in biodosimetry and predictive clinical settings.

To go beyond DEGs, we investigated the transcriptional IR response of whole blood. About 200-300 TFs were commonly regulated by IR independently of the dose and time point. But still, the top-regulated TFs were upregulated depending on the dose. There was a general downregulation of TFs after 6h, especially inflammatory TFs, which might also be attributed to the ‘time in culture’ effect. So far, only Biolatti *et al*. (42) have investigated TFs in a meta-analysis of publicly available microarray datasets of *ex vivo* low linear energy transfer irradiated human whole blood. Their target gene prediction analysis was performed on 275 DEGs revealing TP53, E2F7, NFIA, TCF4, HSF1, JAZF1, KDM4B, and SMPX as overrepresented TFs, partly confirmed by self-performed qPCR. In particular, TCF4 was highlighted as a radiation marker, which showed a general downregulation at all radiation doses examined (0.5-5 Gy). In our study, TCF4 was not found among the DEGs. However, it was downregulated at all doses and time points, except for 0.5 Gy after 2h, which showed a minimal positive log fold change. In this study, the genome-wide transcriptional response in whole blood with or without IR was analyzed for the first time providing important information about these cellular regulatory mechanisms.

When grouping IR doses by their effect on gene expression profiles, we observed that lower doses (≤ 1 Gy) already triggered a significant activation of DDR genes. This activation was evident from 2h post-irradiation and became more pronounced over time. These findings align with the cellular radiation response of the radiosensitive hematologic system, where the DDR can induce apoptosis even at very low radiation doses (∼100 mGy) (8,53–55). At mild levels, these effects contribute to the anti-inflammatory benefits of low-dose RT (e.g., 0.5 Gy) used to treat benign inflammatory degenerative diseases (10). A further potential contributor to the anti-inflammatory effects of low IR doses might be the significant upregulation of CEACAM1, which we observed 2h after exposure to 0.5 Gy. This suggests a dual mechanism involving general DDR activation and specific gene regulation in promoting therapeutic benefits at low radiation doses.

However, IR-induced depletion of leukocytes or even hematopoietic stem cells can have severe consequences, especially during fractionated RT with higher single tumor doses (≥ 2 Gy) or after radiation accidents, resulting in leukopenia or the hematologic acute radiation syndrome (56,57). Beyond hematologic toxicity, IR-associated gene expression signatures in blood are increasingly recognized as predictive markers for individual susceptibility to IR-induced toxicity in solid tissues. Manning *et al*. (19) investigated the expression of 294 inflammation-related genes in the blood of endometrial and head and neck cancer patients during RT. Their study identified consistent regulation of several genes, such as ARG1, BCL2L1, and MYC, during fractionated RT. Additionally, three genes (CD40, OAS2, and CXCR1) correlated with the severity of late normal tissue toxicities. We confirmed IR-induced regulation of some target genes (BCL2L1, MYC, CD40) through *ex vivo* blood exposure to single doses. However, more detailed genome-wide analyses of RT-related gene expression patterns and additional *ex vivo* validation are essential. These efforts could identify predictive biomarkers for the adverse effects of RT and personalization of treatment.

Understanding patient-specific immunological radiation responses is essential, especially in multimodal approaches combining RT with immuno-oncological strategies, such as immune checkpoint inhibitors (13). To maximize synergistic effects, it is crucial to investigate how therapeutic radiation influences the innate and adaptive immune system (3,58,59). Pro-inflammatory reactions are known to be enhanced during RT, with the T-cell response playing a pivotal role in achieving maximum therapeutic efficacy (60). These responses are also triggered by the immunogenic cell death of tumor cells, which release damage-associated molecular patterns activating the DNA-damage-related proinflammatory cGAS-STING-IFN1 signaling pathway (61,62). Our findings confirm that radiation doses ≥ 2 Gy can trigger these effects, even in *ex vivo* irradiated blood samples. Activation of the cGAS-STING-IFN1 pathway was observed at 2 Gy and 4 Gy as early as 2h post-irradiation, though this activation diminished by 6h (Supplementary Fig. S5). The expression of pro-inflammatory cytokines, such as IL1B and TNF, was significantly upregulated in response to IR doses starting at 2 Gy. In contrast, El-Saghire *et al*. (20) have described an IR-induced inflammatory gene signature in human whole blood 8h after *ex vivo* exposure to even a very low dose of 0.05 Gy X-rays. In a subsequent study, the authors showed an inflammatory transcriptomic response involving viral, adaptive, and innate immune signaling 24h after the first 2 Gy fraction of RT in the blood of prostate cancer patients (63). Our previous biodosimetric studies using DNA damage markers revealed peripheral leukocytes receive only about 10% of the tumor dose of 2 Gy during a single session of normofractionated RT (64,65). Only a minor fraction of immune cells residing within the tumor volume or irradiated lymph nodes absorb the full tumor dose per fraction. In line with this and with the data of the present work, recent transcriptome analyses of whole blood from cancer patients during RT showed inflammatory changes at the transcriptome level only at the end of treatment when higher cumulative blood doses were reached (3,16). O’Brien *et al*. (3) identified an IR-specific signature of 15 up- and 16 downregulated inflammatory genes in cancer patients’ blood at the end of RT, independent of the individual immune status and confounding inflammation-related factors. Of these total 31 genes, ALOX5, CEBPB, CD40, CCR7, LTA, and MYC were also detected as DEGs in our analysis. However, the direction of their regulation was partially inconsistent with O’Brien’s results. While fractionated low-dose exposure of peripheral leukocytes likely drives cumulative effects during RT, this aspect of gene expression changes in the blood has not been investigated *ex vivo*. Understanding this repeated low-dose exposure’s impact on leukocyte gene expression could provide insights into systemic immune modulation during RT, patient outcomes, and toxicity. These findings emphasize IR’s ability to trigger pro-inflammatory signaling beyond the tumor microenvironment and highlight the importance of dose-dependent immune effects, especially in radiation-immuno-oncology.

## Conclusion

Our bulk RNA-seq study on *ex vivo* X-ray irradiated human blood identified several novel radiosensitive genes and transcription factors, while also confirming known IR-responsive gene signatures for biodosimetry. The generated dataset enables testing and validating their potential for dose estimation and radiation response prediction *in vivo*. In particular, the transcriptional inflammatory response is of interest for therapeutic IR applications in RT in the context of radiation-immuno-oncology concepts. Our future work will involve extending the current scenario to analyze gene expression dynamics in response to other radiation qualities and dose rates.

## Availability of code, data and materials

All analyses presented in this manuscript were run in R v4.4.1, with a detailed computational workflow and a Snakemake environment definition available athttps://github.com/AhmedSAHassan/Phybion_ShortReads_Analysis. The processed data together with the metadata of the corresponding samples and the analysis code for reproducibility are available on this GitHub repository. The individual-level RNA-seq data (FASTQ format) presented in this article are not publicly available, as they contain sensitive human data underlying data protection rules. Due to the regulations to protect the participants’ data and to ensure that they remain pseudonymized, the raw datasets generated and analyzed in this study are only available upon reasonable request from the corresponding author.

## Acknowledgments

The authors thank Giusy Carlino for her excellent technical support and all donors for providing blood samples. This work has been supported by the computing infrastructure provided by the Core Facility Bioinformatics at the University Medical Center Mainz.

## Authors’ Contribution

Conceptualization: SZ, FM. Data curation: AH. Formal analysis: AH, FM. Funding acquisition: SZ, FM, HS. Investigation: AH. Methodology: FM, AH, SZ, MC, DW. Project administration: SZ. Resources: SZ, FM, HS. Software Supervision: FM. Validation: AH, FM, SZ. Visualization: AH, FM, SZ. Writing – original draft: AH, SZ, FM. Writing – review & editing:: AH, FM, SZ, HS, DW, MC.

## Ethics declarations

The studies involving human participants were reviewed and approved by the Ethics Committee of the Medical Association of Rhineland-Palatinate [No. 2023-17191].

## Supplementary Information

The online version contains supplementary material available at

