## Supplementary figures and images for "Genome-wide transcriptomic response of whole blood to radiation"

### Supplemental Figure 1

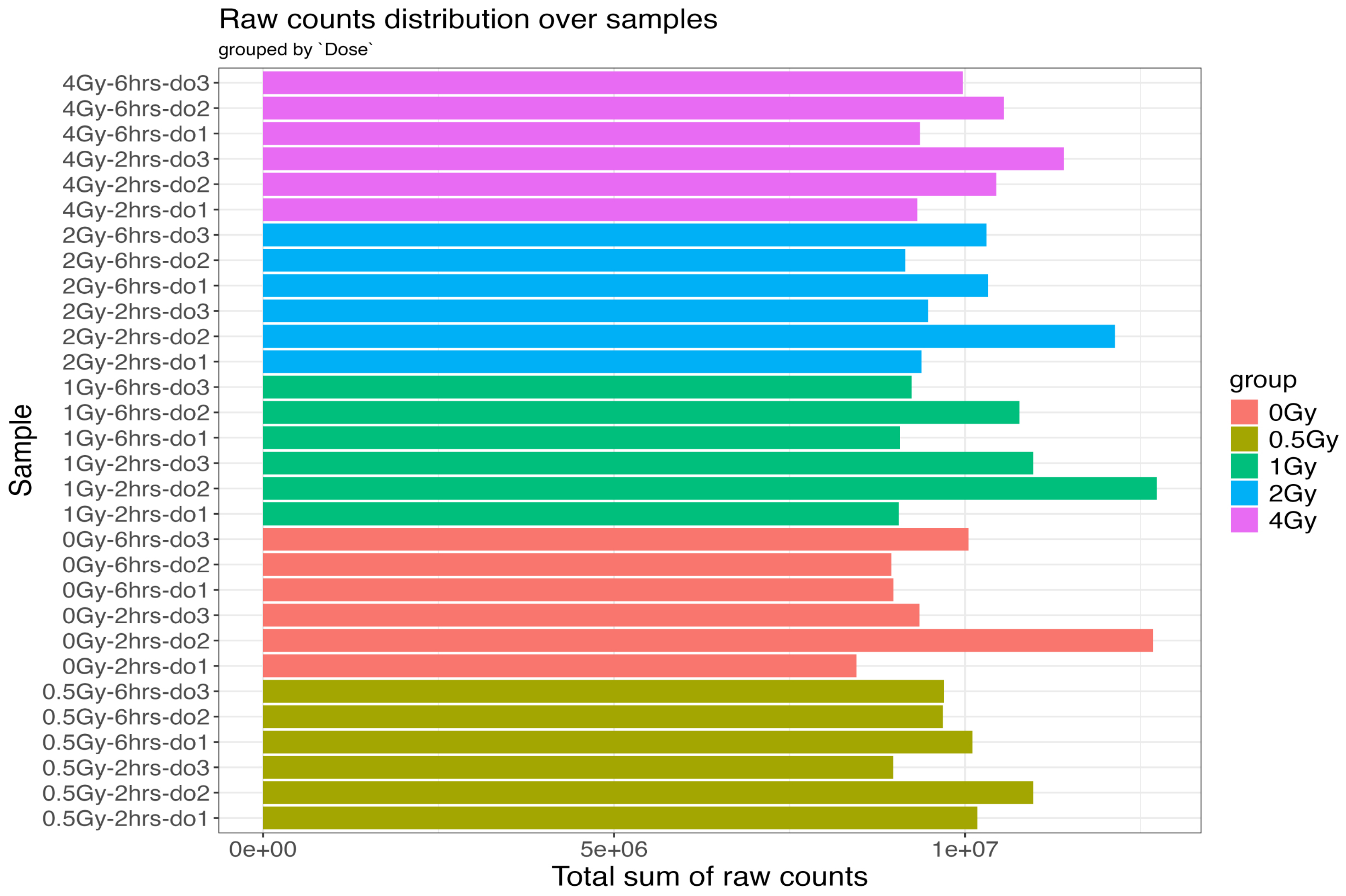


Supplementary Figure S1: Total number of raw reads count associated with each sample.
