## Supplemental Figure 2 for "Genome-wide transcriptomic response of whole blood to radiation"

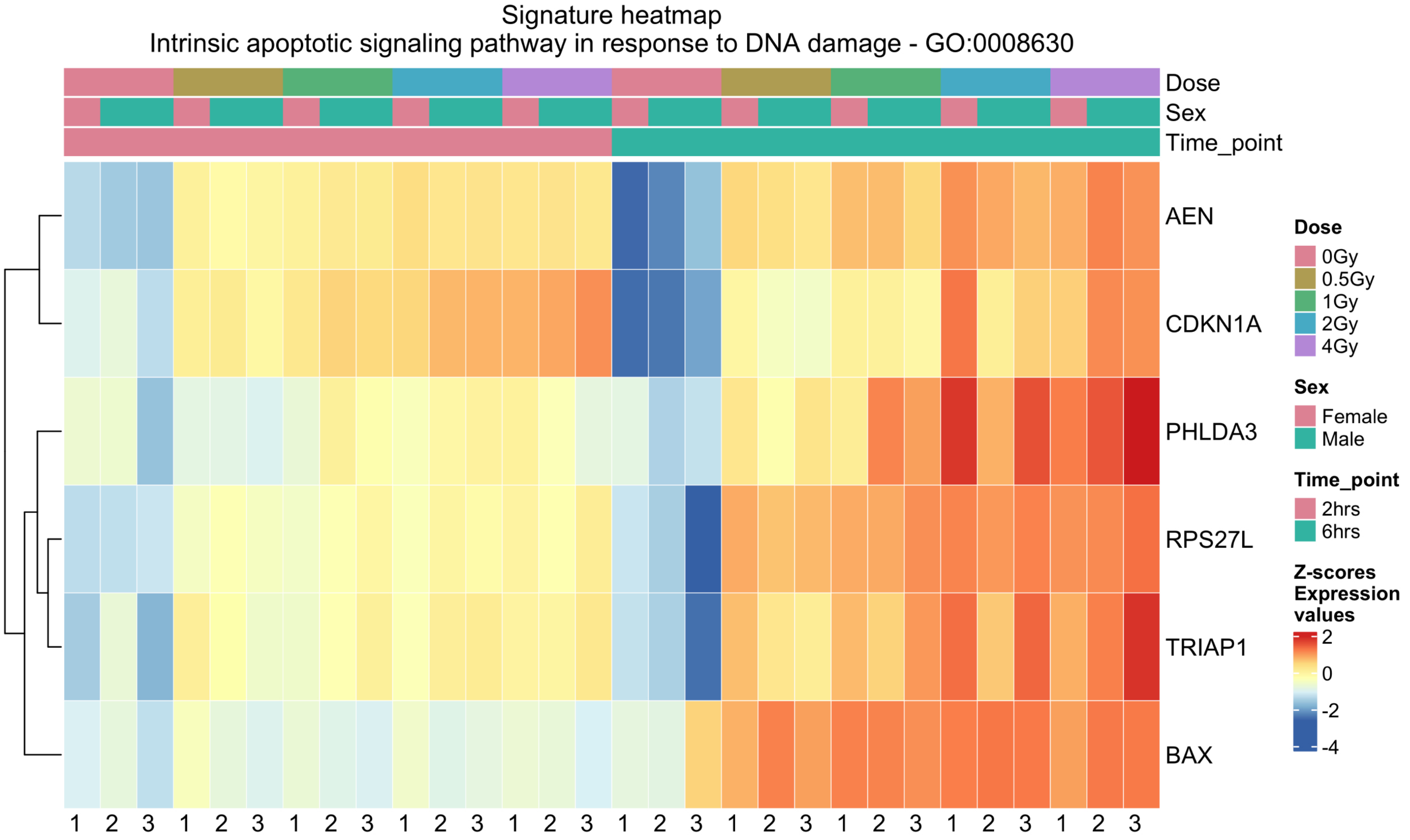


Supplementary Figure S2: Heatmap representing the expression pattern of the differentially expressed genes extracted from 0.5 Gy vs. 0 Gy 6 h post-irradiation associated with the intrinsic apoptotic signaling pathway GO:0008630.
