## Supplemental Figure 3 for "Genome-wide transcriptomic response of whole blood to radiation"

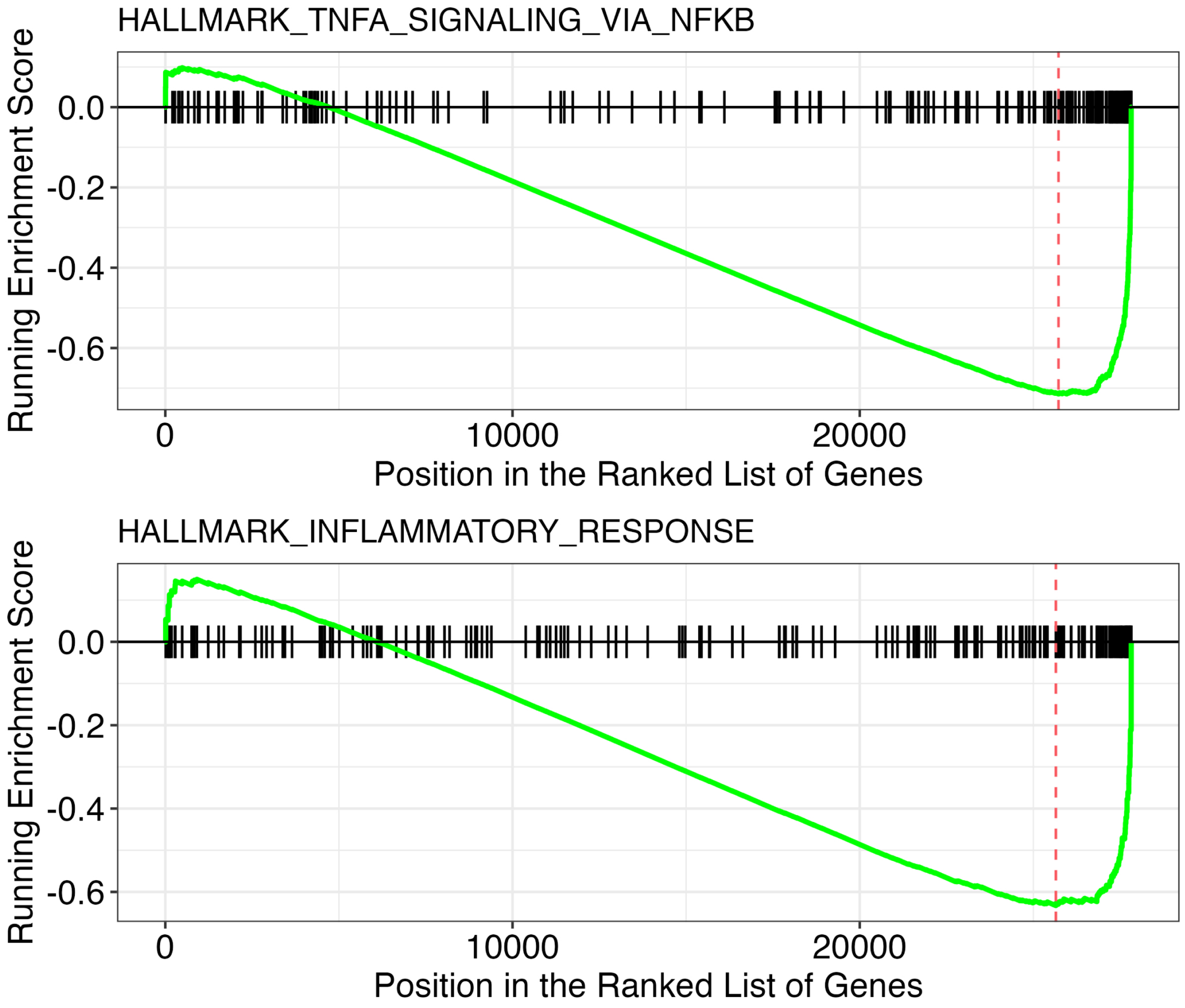


Supplementary Figure S3: Functional annotation by Gene Set Enrichment Analysis with associated normalized enrichment score and p-value. The plot illustrates the transcriptional patters of genes associated with TNF signaling via NFKB (top panel) and inflammatory response (bottom panel) pathways in response to the incubation effect. Genes were ranked by the sign of their LFC multiplied by -log10(pvalue).
