## Supplemental Figure 4 for "Genome-wide transcriptomic response of whole blood to radiation"

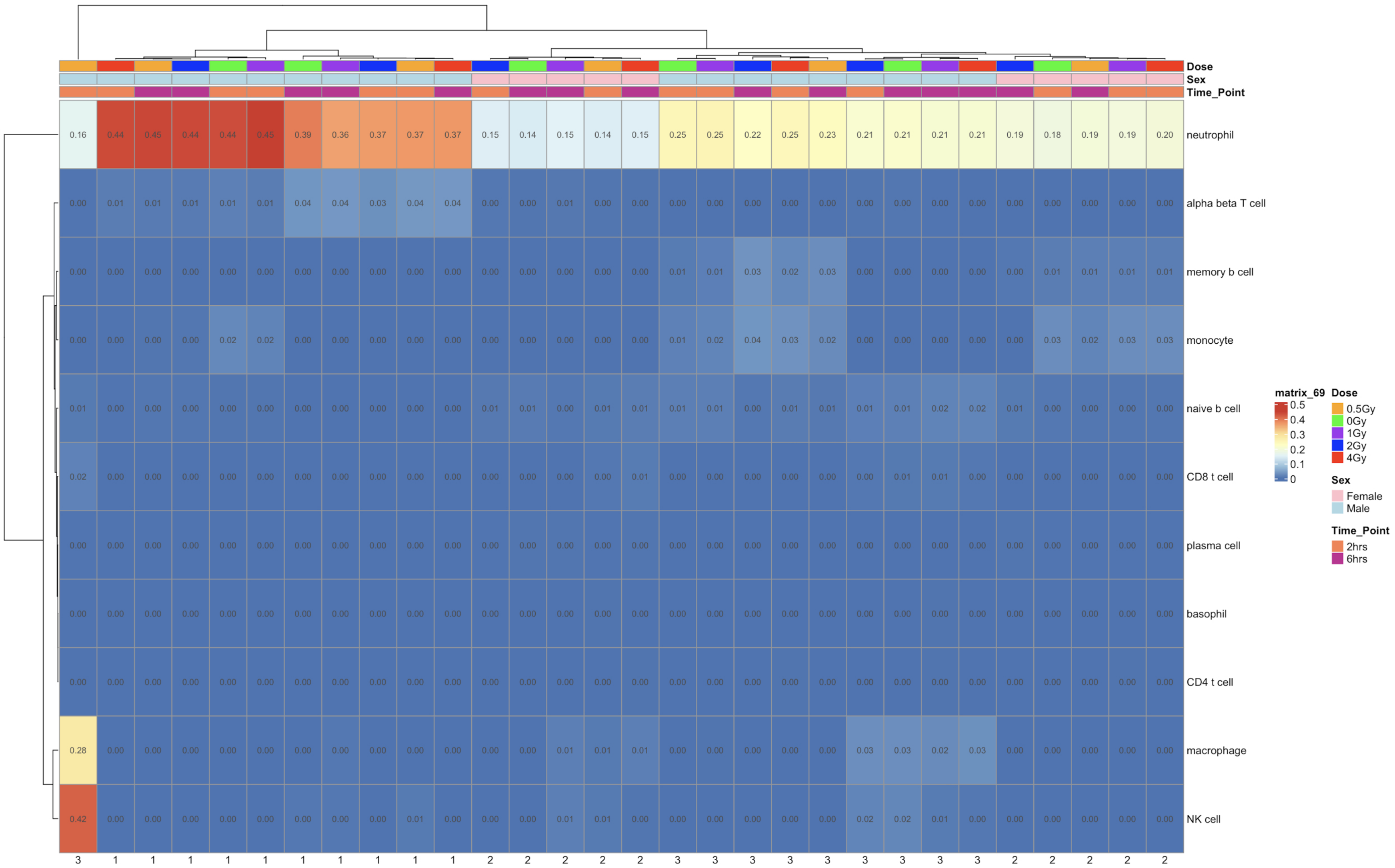


Supplementary Figure S4: Deconvolution heatmap resulting from Omnideconv bayesprism-based estimation of the immune cell type relative abundance.
