## Supplemental Figure 5 for "Genome-wide transcriptomic response of whole blood to radiation"

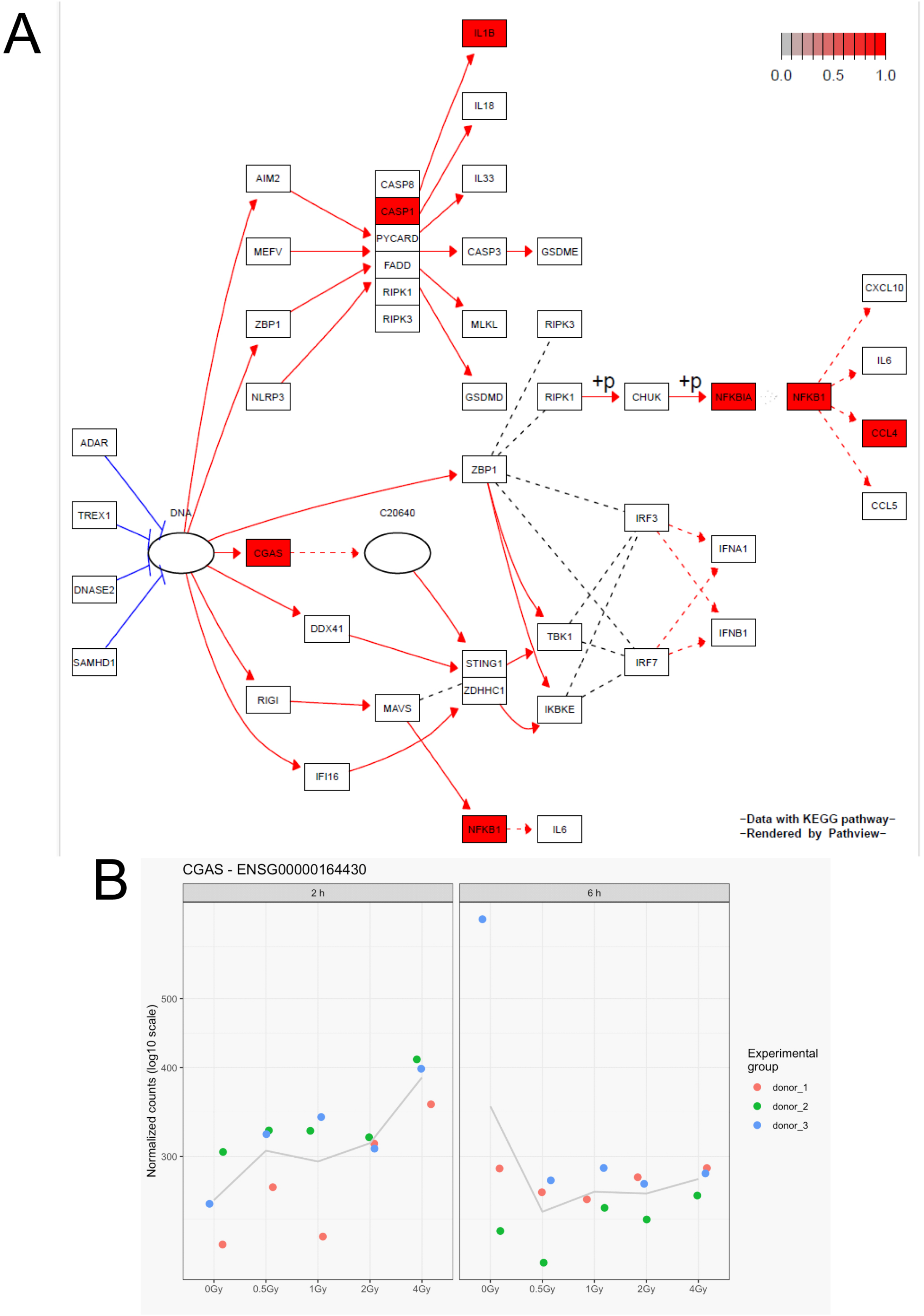


Supplementary Figure S5: (A) cGAS pathway obtained from KEGG (Kyoto Encyclopedia of Genes and Genomes) database and rendered by Pathview. Red-colored genes correspond to differentially expressed genes extracted from 4 Gy vs. 0 Gy comparison 2 h post-irradiation. Solid red arrows represent activation of genes, solid blue lines represents inhibition. +P means phosphorylation. (B) Dose-response relationships of normalized cGAS expression 2 h and 6 h after irradiation.
